# Single-cell characterization of subsolid and solid lesions in the lung adenocarcinoma spectrum

**DOI:** 10.1101/2020.12.25.424416

**Authors:** J. Yanagawa, L.M. Tran, E. Fung, W.D. Wallace, A.E. Prosper, G.A. Fishbein, C. Shea, R. Hong, B. Liu, R. Salehi-Rad, J. Deng, A.C. Gower, J.D. Campbell, S.A. Mazzilli, J. Beane-Ebel, H. Kadara, M.E. Lenburg, A.E. Spira, D.R. Aberle, K. Krysan, S.M. Dubinett

**Author notes:** Current affiliation: Department of Pathology, Keck School of Medicine of USC, Los Angeles, CA 90033. These authors contributed equally (J. Yanagawa, L.M. Tran).

## Abstract

Determining the clinical significance of CT scan-detected subsolid pulmonary nodules requires an understanding of the molecular and cellular features that may foreshadow disease progression. We studied the alterations at the transcriptome level in both immune and non-immune cells, utilizing single-cell RNA sequencing, to compare the microenvironment of subsolid, solid, and non-involved lung tissues from surgical resection specimens. This evaluation of early spectrum lung adenocarcinoma reveals a significant decrease in the cytolytic activities of natural killer and natural killer T cells, accompanied by a reduction of effector T cells as well as an increase of CD4^+^ regulatory T cells in subsolid lesions. Characterization of non-immune cells revealed that both cancer-associated alveolar type 2 cells and fibroblasts contribute to the deregulation of the extracellular matrix, potentially affecting immune infiltration in subsolid lesions through ligand-receptor interactions. These findings suggest a decrement of immune surveillance in subsolid lesions.

## Introduction

Subsolid nodules observed on chest computed tomography (CT) are defined as focal hazy opacifications in which underlying lung structures, such as vessels, remain visible. These may be classified as either pure ground glass nodules (GGNs), or semi-consolidative lesions (dense ground glass), or part-solid nodules (PSNs). PSNs refer to discrete solid components at least 2 mm in diameter that do not correspond to the normal lung anatomy. Subsolid nodules often represent premalignancy or early spectrum lung adenocarcinoma, including premalignant lesions of atypical adenomatous hyperplasia and adenocarcinoma in situ, minimally invasive adenocarcinoma, or lepidic predominant invasive adenocarcinoma (Travis et al., 2016). Studies suggest that subsolid nodules that are predominantly ground glass portend a significantly better prognosis compared to lung adenocarcinomas (ADC) that present as part-solid or solid lesions (Aokage et al., 2018; Berry et al., 2018; Hattori et al., 2017; Kim et al., 2020). Some subsolid nodules remain stable for many years while others may progress to invasive disease. There are no existing biomarkers that accurately predict either lesion progression or regression. This ambiguity interferes with clinical decision-making. For example, questions arise in this context as to whether lung-sparing sublobar resection may be more appropriate than the gold standard of lobectomy for these lesions (Kim and Lee, 2019). This clinical dilemma has become more prominent as the incidence of subsolid nodules has increased due to an increase in both screening and diagnostic chest CT scans.

A greater understanding of the biological determinants of progression to invasive disease is critical to the development and refinement of therapeutic interventions in early spectrum lung ADC. The study of preinvasive lesions is technically challenging because it is difficult to localize and harvest samples for research. These early lesions are often small and exist in the lung periphery, where they are not readily accessible. Some subsolid nodules are ultimately surgically resected, either as the target lesion or as a synchronous lesion within the resected lung. These cases allow acquisition of fresh specimens of early spectrum disease, affording the opportunity to enhance our understanding of the functional molecular and cellular biology of individual subsolid lesions, including the components of the surrounding microenvironment. We present here the results of single-cell RNA sequencing of the continuum of human subsolid lesions ranging from GGNs to PSNs, with a focus on the complex interplay between immune and non-immune cells.

## Results

### Single-cell RNA sequencing reveals differential immune infiltration in pulmonary nodules and associated normal lung tissue

We identified four patients with a singular subsolid nodule and two patients with synchronous subsolid and solid nodules (**Table S1**). Among patients with synchronous subsolid and solid nodules, in one patient (Case 2), the subsolid and solid nodules were in the same lung lobe, and in the other patient (Case 4) two subsolid nodules were present in a different ipsilateral lung lobe. The six patients underwent surgical lobectomy and wedge resection and in total seven subsolid nodules (4 primary and 3 synchronous) and two solid nodules were resected. Pathological evaluation of all nodules revealed the diagnosis of lung ADC. Detailed demographic, pathological, and radiological characteristics are included in **Table S1**.

We performed single-cell transcriptomic profiling of nine nodules (7 subsolid and 2 solid) from six patients along with matching normal lung tissue (distance >2cm from the abnormal lesions) (**Figure 1A**). A total of 88,638 cells passed quality control and were subjected to batch-effect correction before entering the analysis pipeline (see **Methods**). Immune and non-immune cells (**Figure 1B**) were separated based on gene expression of *PTPRC* (CD45) for immune cells, *COL1A1* for fibroblasts, *PECAM1* for endothelial cells, *EPCAM* for epithelial cells, and *SCGB1A1* for secretory cells (**Figure S1A**) and analyzed them independently. Unsupervised graph-based clustering of 60,509 immune cells revealed 30 cell clusters, visualized with the Uniform Manifold Approximation and Projection (UMAP) approach (**Figure S1B**). UMAP plots demonstrated a difference in the spatial distribution of different immune cell types infiltrating normal and abnormal tissues (**Figure 1C**). The thirteen major immune cell types, including natural killer (NK), natural killer T (NKT), CD8^+^ T (CD8), CD4^+^ T (CD4), gamma delta T (γδT), B, plasma, mast, dendritic cells (DC), plasmacytoid DC (pDC), macrophages, monocytes, and neutrophils were identified based on the enrichment scores for the lineage gene sets and the expression of canonical lineage markers (**Figure 1D and S1C**). These cell types were further subjected to sub-lineage analyses to define their functional significance. We then evaluated the frequency of different immune cell types infiltrating normal lung, subsolid, and solid lesions. We observed a significant increase in infiltrating CD4^+^ T, B cells and DCs and reduced NK and NKT cells in subsolid lesions compared to the associated normal tissues (**Figures 1E** and **1F**). These changes were also significant when comparing solid lesions to normal tissue, but not between subsolid and solid lesions (data not shown). The frequencies of adaptive immune cells, including CD4^+^ T, B cells and DCs, were positively correlated (**Figure S1D**). These findings are consistent with other studies (Lambrechts et al., 2018; Lavin et al., 2017) and suggest profound alterations in innate and adaptive components of the immune microenvironment in early spectrum lung cancer.

**Figure 1.**
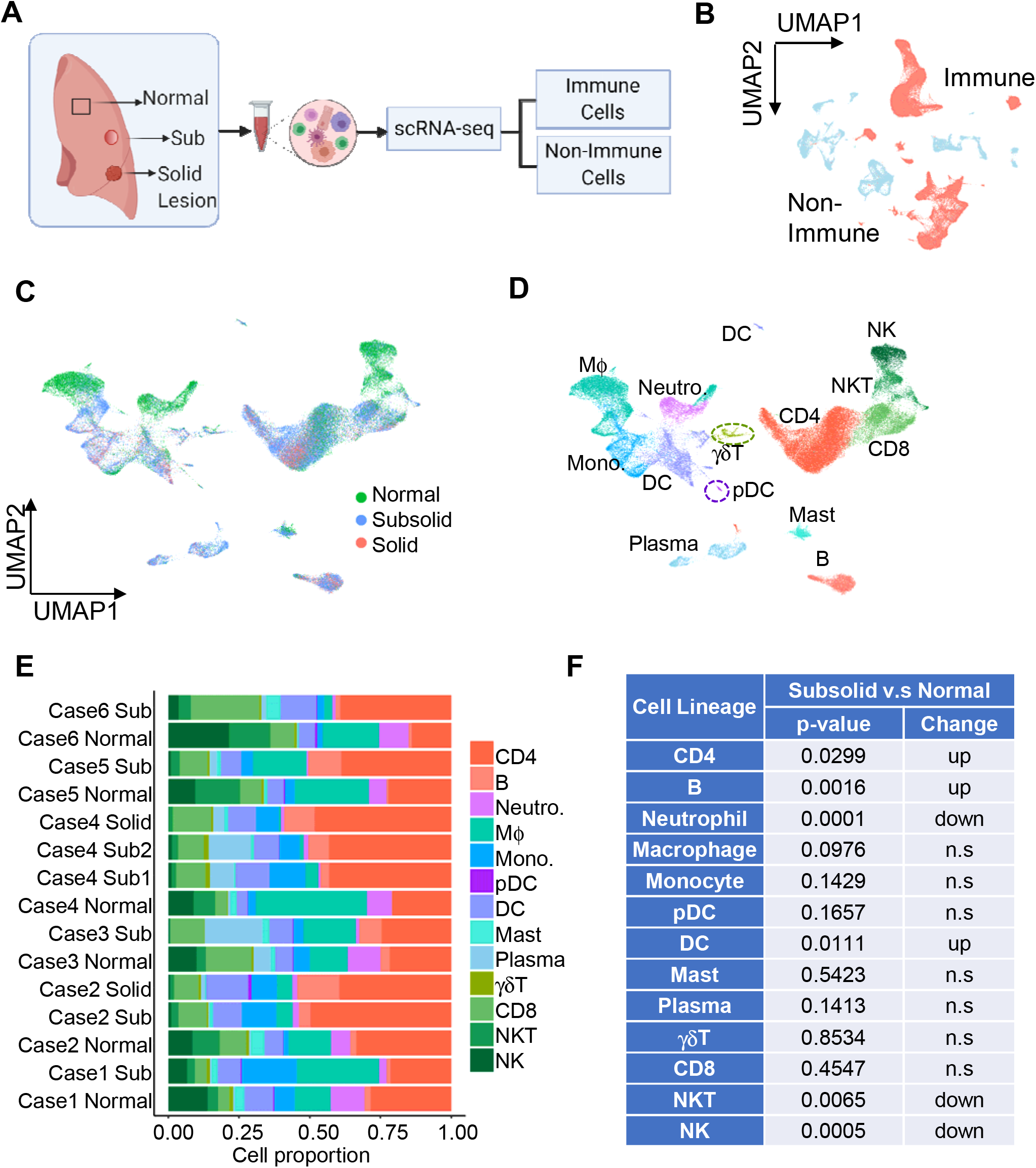
Single-cell transcriptional profile of human immune cells in lung subsolid nodules and associate normal lung tissue. (A) Schematic of the experimental workflow to capture single-cell transcriptome profiles in subsolid nodules, matched non-involved normal lung and solid lesions in six patients (B) Two-dimensional visualization (UMAP plot) of immune (red) and non-immune cells (light blue) (C) UMAP plot visualizing spatial distributions of immune cells based on sample type. Colors represent sample types (D) UMAP plot visualizing thirteen immune cell types. Colors represent cell types, including natural killer (NK), natural killer T (NKT), CD8^+^ T (CD8), CD4^+^ T (CD4), gamma delta T (γδT), B, plasma mast, dendritic cell (DC), plasmacytoid DC (pDC), macrophage (MΦ), monocyte (Mono) and neutrophil (Neutro) (E) The relative proportion to total number of captured immune cells of 13 immune cell types in each sample. Colors represent cell types. (F) Table summarizing the statistical significance between subsolid lesions and normal tissue based on the contribution of each immune cell type. p-values were calculated by the linear mixed effect model.

### Reduction of cytolytic NK and NKT cells in subsolid lesions

There was a profound decrease in the proportion of NK and NKT cells among the infiltrating CD45^+^ cells in all subsolid and solid lesions compared to the associated normal lung (**Figure 1F** and **2A**). Although the numbers of sequenced NKT were similar to those of NK cells, NKT cells formed two distinct clusters, cluster 11 and 20. Cluster 20 was present in approximately 50% of cells originating from malignant lesions, but less than 20% of cells from normal tissue (**Figure 2B**). This suggests that NKT cells infiltrating malignant lesions differed in phenotypic profile from those infiltrating normal lung tissue.

**Figure 2.**
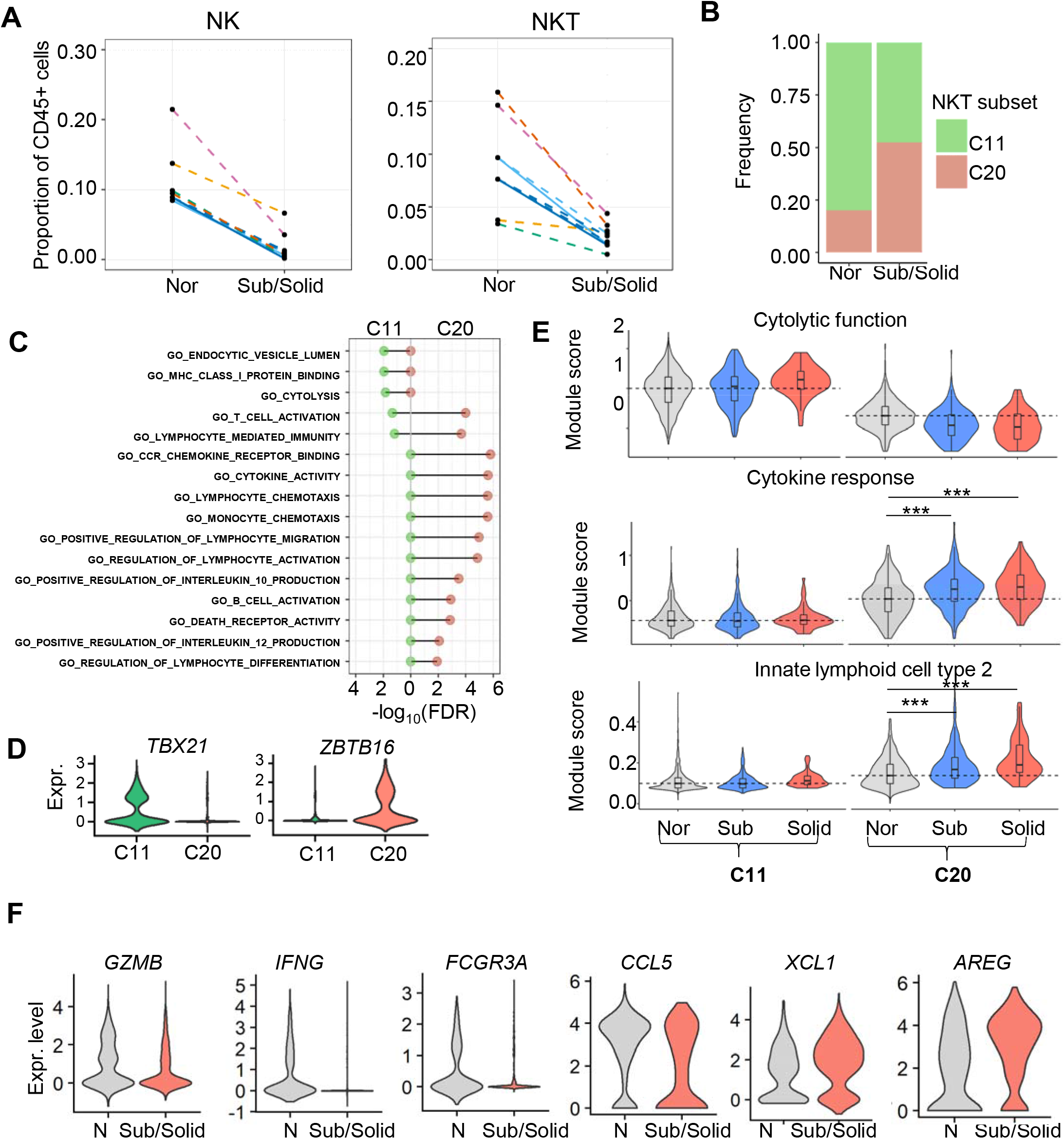
Reduction of cytolytic NK and NKT cells in subsolid lesions. (A) Proportions of immune cells identified as natural killer cells (NK) and natural killer T cells (NKT) in each lesion (dot). Line colors represent individual patients, while line pattern indicates the normal-subsolid (dashed) and normal-tumor (solid) relationship. (B) Distribution of NKT cell subtypes in different lesion types. Colors indicate subtypes (C) Gene Ontology annotation of NKT clusters, C11 (green) and C20 (red) (D) Transcription factor expression associated with invariant NKT subtype 1 (*TBX21*) and 2 (*ZBTB16*) in NKT clusters (E) Violin plots illustrating the distribution of module scores (average z-score of signature genes per cell) in various clusters and lesion types. Dashed lines represent median score based on the normal cells in the selected clusters. ***p<1e-10 based on rank-based Wilcoxon test. (F) DEGs (p<1e-5) assessed by comparing normal lung and subsolid/solid cells in cluster C20 NKT

The biological functions of the NKT clusters were determined by performing gene ontology (GO) analysis. This revealed that gene expression signatures of NKT cells in cluster 11 were enriched in genes associated with cytolytic activity, whereas cluster 20 was enriched in genes encoding cytokines, chemokines, and interleukins involved in modulation of immunocompetent cells, including DCs and T cells (**Figure 2C**). Clusters 11 and 20 demonstrated mutually exclusive expression of transcription factors *TBX21* and *ZBTB16* (**Figure 2D**), which are characteristics of invariant NKT (iNKT) type 1 and 2 subsets, respectively (Terabe and Berzofsky, 2018). To establish an unbiased annotation of NKT clusters as iNKT subtypes, we utilized the gene modules derived from either NK cells or innate lymphoid cells (ILC) based on the premise that iNKT subtype functions are similar to their NK and ILCs counterparts(Bennstein, 2017; Terabe and Berzofsky, 2018). Crinier and colleagues have identified two transcriptome signatures conserved across organs and species, encompassing NK1 and NK2 signatures associated with cytolytic activity and cytokine responses, respectively (Crinier et al., 2018). Calculation of the module score, the average of z-scores across genes defined by the module, revealed highly expressed NK1 signatures in cluster 11 and NK2 signatures in cluster 20, respectively (**Figure 2E**). Within cluster 20, both subsolid and solid cells expressed NK2 signatures that were significantly higher than their normal-derived counterparts (Wilcoxon’s rank test p-value < 1e-10). Signature genes derived from studying murine ILC subsets (Robinette et al., 2015) also defined cluster 20 as ILC type 2 (**Figure 2E**). These data indicate that cluster 11 and cluster 20 represent iNKT1 and iNKT2 cells, respectively. The analysis of the top differentially expressed genes (DEGs) among cluster 20 NKT cells revealed an absence of cytolytic markers (*GZMB, IFNG*, and *FCGR3A/CD16*) and a decrease of *CCL5*, which has T lymphocyte recruiting capacity, in cells derived from the subsolid and solid lesions (**Figure 2F** and **Table S2**). The NKT cell cluster 20 in subsolid and solid lesions was enriched for the DC recruiting chemokines *XCL1* and *AREG* (**Figure 2F**).

### Modulation of adaptive immune responses in subsolid lesions

The adaptive immune cell profiles were assessed in subsolid and solid lesions. Among T cells, a total of 20,445 CD4^+^ cells 5,790 CD8^+^ cells, and 531 γδT cells were identified based on their lineage markers (**Figure S1C and S2A**). Among the infiltrating CD45^+^ cells, the proportions of CD4^+^ T cells were higher in the majority of the subsolid and solid lesions compared to the associated normal lung (**Figure 1F** and **3A**). Only one subsolid lesion (Case 1) had a lower percentage of CD4^+^ T cells compared to the matching normal tissue. The CD4^+^ T cells were aggregated into five clusters and assigned to four subsets, including effector memory T cells (Tem), central-like memory T cells (Tcm), regulatory T cells (Treg), and activated T cells (Tact), based on their canonical markers (**Figure 3B-C** and **S2A**). Compared to other CD4^+^ T cell clusters, the regulatory T cells (cluster 7) expressed elevated levels of regulatory markers, including *FOXP3, IL2RA (CD25)*, and *TNFRSF1* (**Figure 3C** and **S2A**). Although all other CD4^+^ T cell clusters also expressed activation marker *CD44*, they highly expressed tissue-resident marker *CD69*, and two memory markers (*IL7R/CD127* and *PRDM1/BLIMP1*). The Tem cell cluster (cluster 1) was distinguished from other CD4^+^ T cell clusters by expression of *IFNG, XCL1*, and genes encoding granzymes, while the Tcm cell cluster (cluster 0) was identified based on its markers *CCR7* and *TCF7*. Activated T cells (clusters 14 and 21) expressed lower levels of *CD44* and *IL7R* compared to Tem and Tcm cells, but had comparable levels of *CD69*.

**Figure 3.**
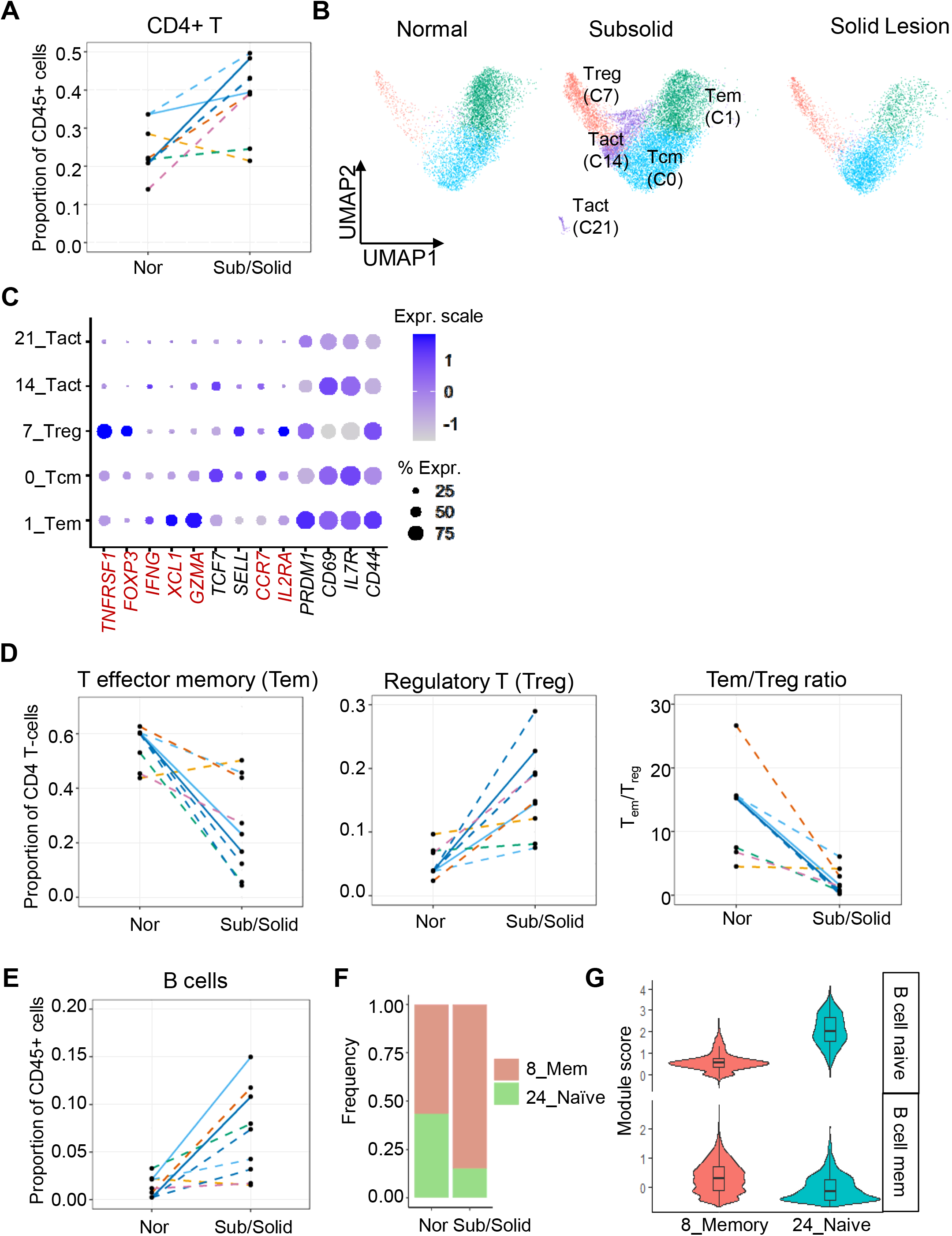
Profiles of the adaptive immune populations in subsolid nodules. (A) CD4^+^ T cell proportions among immune cells in each sample (dot). Line colors represent individual patients, while line pattern indicates the normal-subsolid (dashed) and normal-tumor (solid) relationship. (B) UMAP plots illustrating the distribution of CD4^+^ T cell subsets in different tissues. Colors indicate CD4^+^ T cell sub-lineages, including effector memory T cells (Tem), central-like memory T cells (Tcm), regulatory T cells (Treg), and activated T cells (Tact) (C) CD4^+^ T cell sub-lineage expression makers in individual clusters illustrated by bubble plots. Functional genes in red were also identified as the cluster markers (D) The proportion of CD4^+^ T cells identified as Tem (left) and Treg cells (middle), as well as their Tem:Treg ratio (right) in each sample (dot). Line colors represent individual patients, while line type indicated the normal-subsolid (dashed) and normal-tumor (solid) relationship (E) B cell lineage frequency in each lesion (F) Distribution of B cell subtypes in different lesion types. Colors indicate subtypes (G) Violin plot illustrating the distribution of module scores associated with B cell functions.

Analysis of CD4^+^ T cell subset frequencies revealed differences among normal lung, subsolid and solid lesions (**Figure 3D**). The proportion of Tem among infiltrating CD4^+^ T cells decreased significantly (mixed linear model p-value = 0.016), while the proportion of Treg increased (mixed linear model p-value = 0.015) in subsolid nodules compared to the matching normal tissue (**Figure 3D – left** and **middle panels**). The Tem:Treg ratios, a measure of T-cell mediated immune activity, was decreased in the subsolid lesions (**Figure 3D – right panel**). These results suggest that the increase of CD4^+^ T cells infiltrating subsolid lesions may be indicative of immune suppression rather than effective immune surveillance.

The characterization of infiltrating CD8^+^ T cells in subsolid and solid lesions revealed no significant differences in the proportion of CD8^+^ T cells to total immune cells between malignant lesions and normal lung (**Figure 1F and S2B**). Based on lineage markers, CD8^+^ T cells formed two clusters associated with exhausted (cluster 5) and activated (cluster 6) subsets (**Figure S2A**). There was no significant difference in the percentages of CD8+ T cell subsets between malignant lesions and normal tissue (**Figure S2C**). As determined by DEG analysis, the most down-regulated genes in tumor-infiltrating activated CD8^+^ cells were those associated with cytotoxic function (*GZMB* and *GNLY*) and the movement of molecules across the nuclear envelope (*LMNA*) (**Figure S2D**).

An increase in B cell infiltration was observed in subsolid lesions compared to normal lung tissue (**Figure 1F** and **3E**). Graph-based clustering analysis revealed that B cells existed in two subpopulations: large (cluster 8) and small (cluster 24) groups, which were present in different proportions between malignant lesions and normal lung tissue (**Figure 3F**). The proportions of cluster 8 were 0.83 and 0.47 among B cells originating from malignant lesions and normal tissue, respectively. Analysis of B cell lineage markers (Newman et al., 2015) revealed that B cells in cluster 24 highly expressed naïve markers, while those in cluster 8 highly expressed memory markers (**Figure 3G**), suggesting that the majority of infiltrating B cells mediate memory functions.

### Modulation of DC and monocyte phenotypes in subsolid lesions

The increase in CD4^+^ T and B cells in malignant lesions was positively associated with DC infiltration that was also significantly increased in malignant compared to normal lung tissue (**Figure S1D**). 5,363 DCs and 185 plasmacytoid DCs (pDCs) (cluster 28) were identified. Graph-based clustering produced five DC clusters. Two large groups were composed of approximately 2000 cells per cluster (cluster 10 and 12), and three minor groups were composed of several hundred cells per cluster (cluster 23, 26, and 27). DC groups were characterized utilizing gene modules (Villani et al., 2017) in which DCs were classified into six subtypes: DC1 (conventional DC1 or cDC1); DC2 (conventional DC2 or cDC2); DC3 expressing CD14^+^ monocyte markers in addition to cDC2 markers; DC4 expressing CD16^+^ monocyte markers; rare DC5, and DC6 as pDC subtypes. The smallest, cluster 27, expressed the DC5 gene module, the second smallest, cluster 26, expressed DC1/cDC1 markers, while the largest, cluster 10, expressed cDC2 markers (**Figure S2E**). The lineages of clusters 10 and 26 were confirmed by their high expression of lineage markers CD1C and CADM1, respectively (**Figure S1C**). Cluster 12, the second-largest, highly expressed markers associated with monocyte-derived DC3 and DC4, and moderately expressed cDC2 markers, suggesting that cluster 12 is comparable to the DC3 subtype. Cluster 23, a small cluster, expressed markers associated with monocyte-derived DC3 and DC4, but expressed DC6/pDC rather than a cDC2 lineage. Cluster 23 was one of two clusters that highly expressed proliferation genes, including *MKI67* (**Figure S1C**). The results suggest that cluster 23 cells may represent a transition state in which DCs differentiate from monocytes. Among DCs, the proportion of cDC2 substantially increased, while that of cDC1 decreased in subsolid and solid lesions compared to normal tissue (**Figure S2F**). cDC1 and cDC2 mediate antigen recognition in CD8^+^ and CD4^+^ T cells through MHC class I- and II presentation, respectively. As expected, the percentages of infiltrating CD4^+^ T cells were more highly associated with cDC2 than to total DCs (Pearson correlation coefficient 0.78 vs. 0.65). In contrast, the correlation between infiltrating CD8^+^ and cDC1s was limited. Compared to the normal lung, the cDC1:cDC2 ratio was decreased (mixed linear model p value = 0.06) in subsolid lesions, with the exception of Case 4 lesion 2 and Case 5 (**Figure S2G**), suggesting a decrement of CD8^+^ T cell-mediated immune surveillance activities.

Monocytes (n = 4084 cells) were characterized into subtypes based on monocyte gene modules (Villani et al., 2017). There were no significant differences in infiltration levels between normal lung and associated malignant lesions (**Figure 1F**). Two clusters were associated with monocytes (**Figure S2H**). The large monocyte group (cluster 3) comprised more than 75% of monocytes originating from malignant lesions, and they highly expressed a non-classical monocyte type 2 (CD14+CD16++) signature (**Figure S2H-I**). The minor cluster (cluster 18) had a classical monocyte (CD14++CD16-) signature and were found in 50% of normal tissue-derived monocytes. This suggests that subsolid lesions were enriched by non-classical monocyte type 2 cells.

### Decreased expression of differentiation markers in subsolid lesion alveolar type 2 pneumocytes

Clustering of non-immune cells (28,129) revealed 34 clusters, which were assigned to different cell types through enrichment analysis and canonical marker expression, as described in **Methods** (**Figure S3A-C**). The following cell types were identified: fibroblasts (expression of *COLA1A*), endothelial cells (*PECAM1*), and five different airway lineages, including alveolar type 1 (AT1; *PDPN*), alveolar type 2 (AT2; *EPCAM* and *NKX2-1*), Clara (*SCGB1A1* and *SCGB3A1*), bronchioalveolar stem cells (BASC; *SFTPC* and *SCGB1A1*), and ciliated cells (*FOXJ1*). Many AT2 and BASC clusters were sample-specific, while clusters from other cell types were present in multiple samples (**Figure S3C, lower panel**). In sample-associated AT2 clusters, expression of surfactant encoding genes (*SFTPC, SFTPA1*) was downregulated compared to the non-sample specific counterparts. These AT2 clusters were enriched with cells from subsolid and solid lesions (**Figure S3D**). By pooling AT2 clusters together based on *SFTPC* expression level, two major AT2 subtypes associated with malignant and normal tissues were obtained (**Figure 4A-C**). Malignant-associated AT2 cells had decreased expression of surfactant encoding genes but continued to express other AT2 markers, therefore, the module scores were still higher than those in the other cell types (**Figure 4D**). These results imply the presence of high gene expression heterogeneity in malignant AT2 cells.

**Figure 4.**
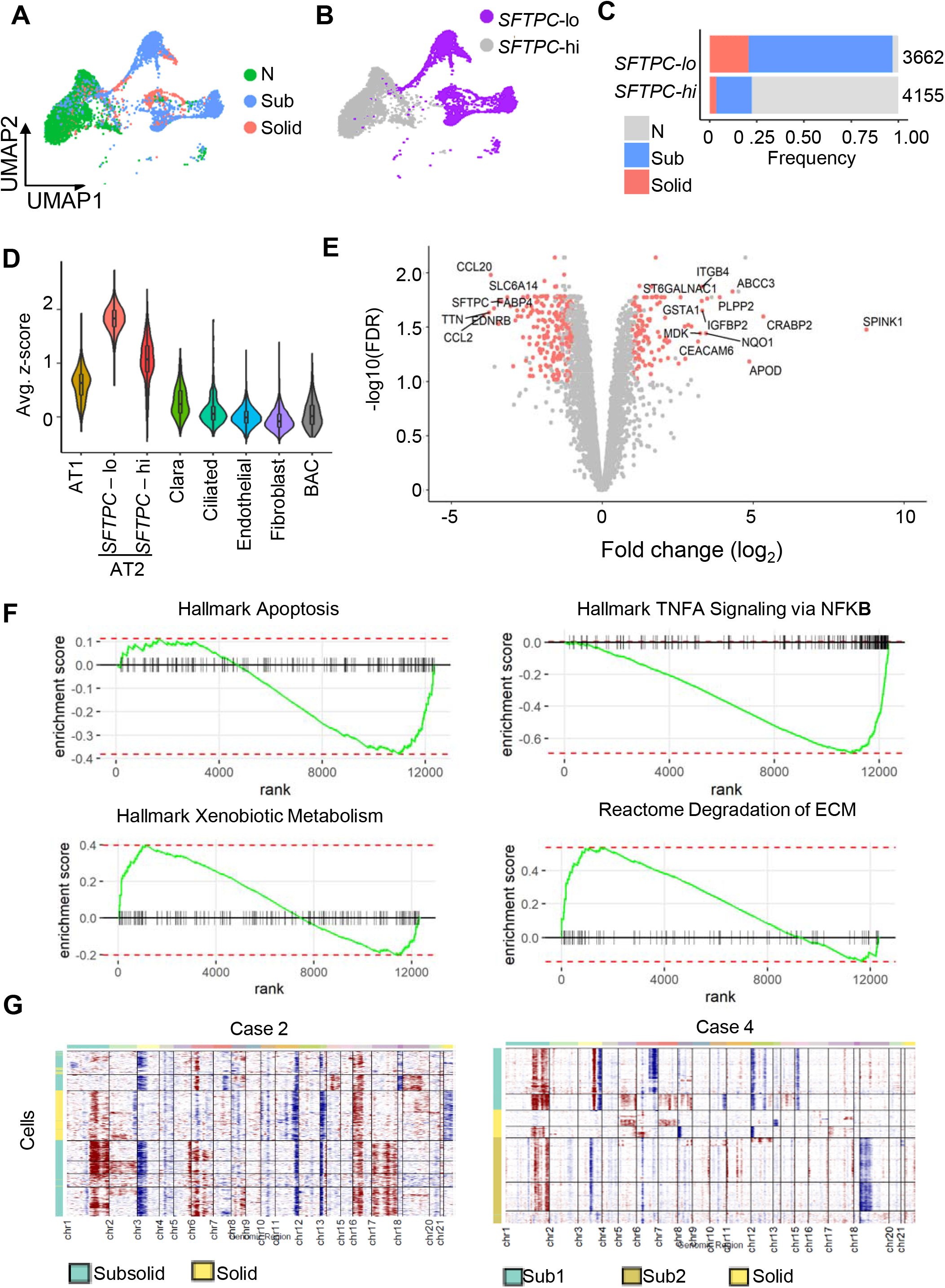
Deregulation of AT2 cells in subsolid and solid lesions. (A) UMAP plot illustrating AT2 cells from subsolid and solid lesion aggregated to clusters distinguished from normal lung tissue. Color represents lesion types (B) Two AT2 clusters identified based on *SFTPC* expression. Colors represent groups (C) Lesion components in two *SFTPC*-based groups. Values indicate cell numbers in the associated groups. Colors indicate lesion types. (D) Violin plots of module scores representing AT2 lineage marker expression in various non-immune populations. (E) Volcano plot representing DEGs in AT2 cells by comparing malignant and normal samples. The p-value (y-axis) and fold changes (x-axis) were determined by utilizing the edgeR approach. The red dots indicate DEGs identified by both edgeR and single-cell modeling MAST approach (F) GSEA plots of the top deregulated pathways in subsolid/solid AT2 cells compared to normal lung cells (G) Heatmaps demonstrating inferred CNAs of malignant (i.e. low *SFTPC* expression) AT2 cells (row) in two subjects with multiple lesions sequenced

Due to this heterogeneity, both single-cell and pseudo-bulk models were utilized to determine the DEGs in AT2 cells (See **Methods**). DEGs (n = 372) were identified between malignant- and normal-derived AT2 cells (**Figure 4E**, **Table S3**). In addition to *SFTPC*, the most down-regulated genes in malignant-derived AT2 cells were *EDNRB* and *TNN*. Downregulation of *EDNRB*, encoding a nonselective endothelin B receptor that induces apoptosis upon activation, has been associated with poor prognosis in NSCLC (Wei et al., 2020). *EDNRB* downregulation may be due to hypermethylation of its promoter region (Knight et al., 2009). The top-upregulated genes, including *SPINK1, CEACAM6, IGFBP2*, and *ABCC3*, are known for their tumorigenic roles in NSCLC and other cancers (Blumenthal et al., 2007; Li et al., 2020; Mehner and Radisky, 2019; Zhao et al., 2013). For example, *SPINK1* is a secreted protein overexpressed in multiple cancers and promotes tumor progression by altering the tumor microenvironment (Chen et al., 2018). It is also positively correlated with poor prognosis in NSCLC (Guo et al., 2019). Pathway analysis indicates down-regulation of the apoptosis and NFKB pathways as well as upregulation of genes regulating extracellular matrix (ECM) degradation and drug metabolisms in malignant-derived AT2 cells (**Figure 4F**).

Copy number alterations (CNAs) were assessed in malignant-associated AT2 cells characterized based on low *SFTPC* expression utilizing a mRNA-inferring CNV approach (Tickle et al., 2019). The analysis was applied to two cases with multiple lesions sequenced in order to evaluate sub-clonal heterogeneity. More than one sub-clone was observed in each sample (**Figure 4G**). In Case 2, cells in the subsolid and solid lesions shared an arm-level amplification of chr16p. The Case 2 samples also had deletions in Chr11q and amplification in Chr1q although the breakpoint was not the same. In addition to sharing the recurrent CNA regions, each lesion harbored distinct alterations, such as Chr3p deletions and Chr6p amplifications in the subsolid lesion, and arm-level deletions of Chr21 in the solid lesion. The genetic tests indicated that the lesions had different *KRAS* genotypes (**Table S1**), suggesting these lesions might have the same ancestry with Chr16p deletion clonal, but developed different subclones independently during the malignant progression. Two subsolid lesions in the right middle lobe and one solid lesion in the right lower lobe were sequenced in Case 4. Most cells in subsolid lesions shared an amplification in Chr1q while those in the solid lesion also carried Chr1q amplification but at a different break point. These three lesions did not have clonal CNA and harbored different driver mutations (**Table S1**), implying that the subsolid and solid lesions originated from different ancestry. Some cells in the subsolid lesion 1 and solid lesion shared arm-level gains of Chr5 and 7, which was absent in the associated less invasive subsolid lesion 2. These gains of Chr5 and 7 were frequently detected in the TCGA lung adenocarcinoma cohort (Cancer Genome Atlas Research, 2014), suggesting these CNAs may contribute to aggressive tumor behavior. The results from CNA inference suggest that the subsolid and solid lesions were genetically heterogenous with multiple sub-clones that evolved independently though possibly originating from the same ancestry.

### Cancer-associated endothelial cells and fibroblasts modify the subsolid nodule microenvironment

Endothelial cells and fibroblasts contribute to the hallmarks of cancer. Nine clusters composed of 8,280 cells, annotated as endothelial cells (ECs), were identified based on enrichment analysis and high *PECAM1* expression (**Figure S3A-C**). Among them, cluster 7 was strongly associated with a specific sample, while the remainder of the clusters were present in multiple samples (**Figure S3C-lower panel**). The majority of these cluster cells expressed the capillary marker *CA4*, while others expressed markers associated with arterial, venous, and lymphatic ECs (**Figure S4A**). Clusters were characterized based on the average expression of EC signature gene modules (e.g., capillary, arterial, venous, immature stalk, and lymphatic ECs), rather than relying on single markers, as described by Goveia et al. (Goveia et al., 2020). Five of nine clusters were identified as capillary ECs, while the remainder ECs were arterial, venous, immature, and lymphatic (**Figure S4B**). ECs from solid lesions were found only in clusters annotated as venous (cluster 14), immature (cluster 15), and lymphatic (cluster 31) EC. Two clusters (cluster 14 and 15) were significantly enriched in subsolid-derived ECs (**Figure 5A-B** and **S4B**). In addition to expressing immature stalk gene signatures associated with the proliferation of ECs, cells in these two clusters also highly expressed markers associated with tip cells that are responsible for EC migration (**Figure 5C**). Both immature stalk and tip-like ECs are building blocks for sprouting angiogenesis. **Figure 5D** illustrates expression of the cluster’s top markers associated with endothelial stalk cells (*ACKR1*) (Chen et al., 2019), tip cells (*INSR, LAMA4*) (Goveia et al., 2020; Stenzel et al., 2011) and increased basal permeability in the blood-brain barrier (*PLVAP*) (Wisniewska-Kruk et al., 2016). These findings indicate an increased potential for angiogenesis in subsolid lesions.

**Figure 5.**
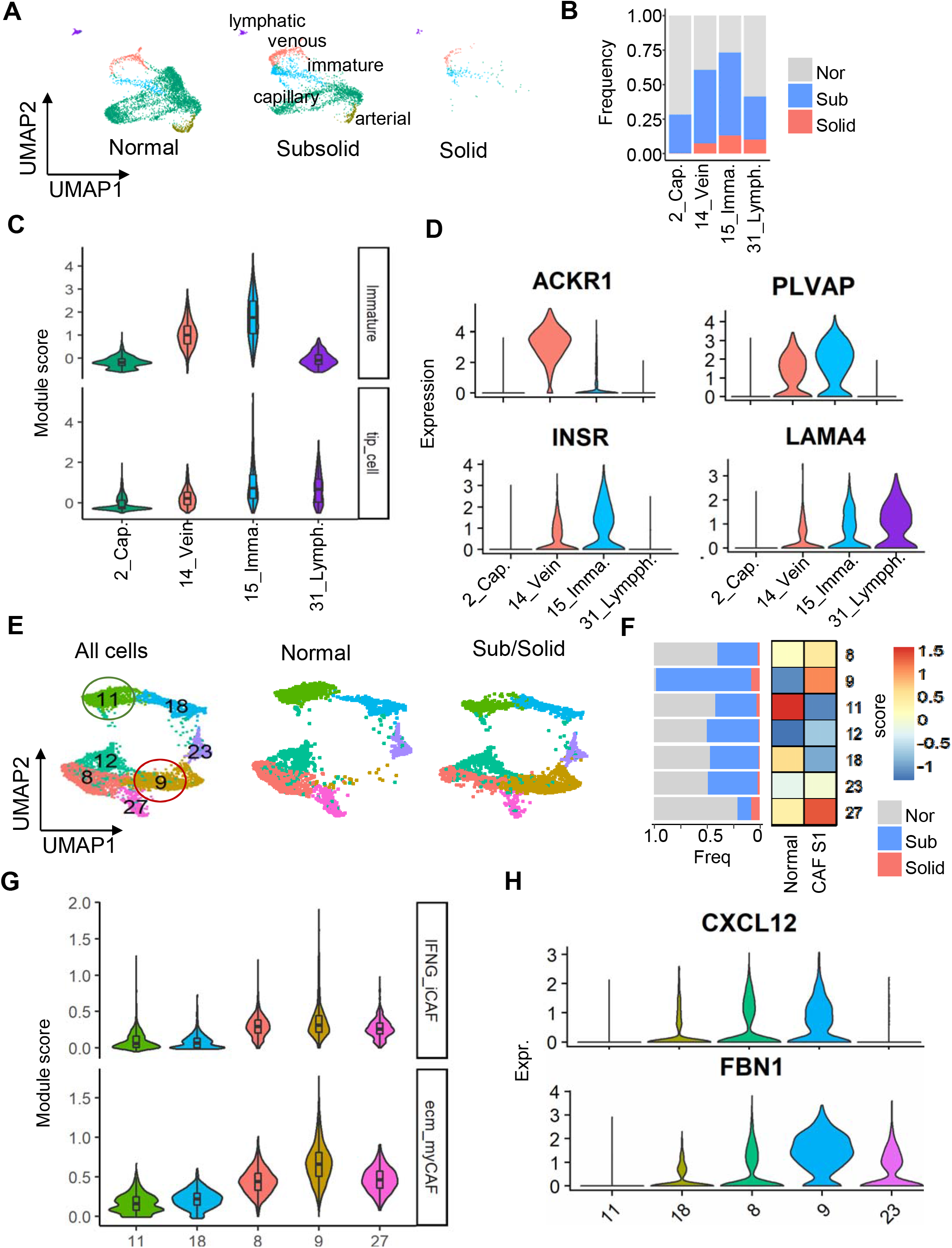
Cancer-associated endothelial cells (ECs) and fibroblasts (CAFs) enriched in subsolid lesions. (A) UMAP plot visualizing the distribution of five EC subtypes in normal lung (left), subsolid (middle), and solid (right) (B) Lesion-based contributions (left) to four clusters representing EC subtypes enriched by either normal lung or subsolid/solid lesions. Color indicates lesion type. (C) Violin plots of module scores associated with cancer-associated immature and tip-like ECs in representative EC clusters. (D) Expression of top DEGs in the representative clusters. DEGs were assessed based on comparing ECs between subsolid/solid lesions and non-involved tissue. (E) UMAP plot visualizing seven fibroblast clusters in all tissue (left) or separated tissues, including normal lung (middle), and subsolid/solid (right) (F) Fibroblast clusters characterized by lesion-based contributions (left) and module scores associated with normal fibroblasts and CAFs (right). Horizontal bar plot (left) indicates tissue proportions present in each cluster. Heatmap (right) represents scores of the normal and CAF gene modules (columns) in clusters (rows). (G) Score distribution of gene modules associated with CAF subtypes in selected clusters representing normal fibroblasts (C11 and 18), CAFs (C8 and 9), and a mixed population (C27). (H) Expression of signatures associated with ecm-myCAFs in the representative clusters

The fibroblast population of 5,886 cells clustered into seven clusters (**Figure 5E**). While most clusters tended to be comprised of an equal mixture of cells derived from both malignant and normal tissues, cluster 9 was predominantly composed of cells derived from malignant samples (**Figure 5E-F**) and expressed canonical cancer-associated fibroblast markers *FAP, S100A4*, and *PDGFRA* (**Figure S4C**). Functional analysis using gene modules associated with normal- and cancer-derived fibroblasts (Kieffer et al., 2020) confirmed that cluster 9 cells expressed CAF-S1 (FAP^hi^SMA^hi^CD29^med-hi^) signatures and identified clusters 11 and 18 as normal fibroblasts (**Figure 5F**). Clusters 8 and 27, neighbors to cluster 9, also expressed CAF signatures but at lower levels. The remaining clusters (12 and 23) displayed mixed signals of normal-derived fibroblasts and CAFs, which illustrates the plasticity in fibroblast transcriptome profiles. CAF subpopulations were further characterized utilizing the gene markers associated with eight CAF subtypes, recently identified in a breast cancer study (Kieffer et al., 2020). In cluster 9, the highest expressed gene modules were associated with extracellular matrix remodeling myofribroblasts (ecm-myCAF) and IFNγ responding inflammatory CAF (IFNγ-iCAF) (**Figure 5G**). Other CAF clusters, 8 and 27, also significantly expressed these gene modules compared to normal fibroblast clusters (11 and 18) but were specific for inflammatory CAFs responding to IL (IL-iCAF) and detoxification (detox-iCAF), respectively (**Figure S4D**). Analysis of the cluster markers revealed collagen encoding genes (*COL1A1, COL3A1, COL5A1*, and *COL6A1*) as the highest expressed markers in cluster 9. Cluster 9 cells significantly expressed *CXCL12* and *FBN1* markers of IFNγ-iCAF and ecm-myCAF (**Figure 5H**). These results suggest that ecm-myCAF and IFNγ-iCAF may play roles in modifying the tumor microenvironment which, in turn, may modulate immune cell infiltration.

### Ligand-receptor interactions in the developing tumor microenvironment

To explain alterations in the immune contexture observed in subsolid and solid lesions compared to the normal tissue, relationships among cellular phenotypes were investigated through their ligand-receptor (LR) interactions. This LR assessment is based on the premise that the upregulation of ligand expression in cell A can either induce expression of the related receptor in cell B or induce migration of cell B to the local region of cell A. The LR interaction strength in each sample was defined as the product of ligand and receptor expressions in related cells, as suggested by Kumar et al (Kumar et al., 2018), and then utilized to determine if the interaction was significantly increased in subsolid and solid lesions compared to normal lung tissue (see **Methods**). LR interactions among AT2, ECs, and CAFs were found to be robust in the subsolid and solid lesions (**Figure 6A**). The CAFs interacted with the malignant-derived AT2 cells and ECs, predominantly by ligands expressed in CAFs (source cells) and receptors expressed by the other two cell types (receivers), rather than vice versa. The interaction between AT2 and endothelial cells in subsolid lesions was found to be a mutual relationship in which each acted equally as source and receiver cells. In the subsolid and solid lesions, the non-immune to non-immune interactions were higher than those of immune to non-immune interactions. These interactions maintained a specific direction in which non-immune cells predominantly acted as sources while immune cells acted as receivers (**Figure 6B**). The non-immune cells frequently communicated with Treg cells, B cells, DCs, and monocytes (**Figure 6B-left**), suggesting that changes in transcriptome profiles of non-immune cells may determine the immune contexture of the microenvironment. Among immune cells, only DCs and monocytes acted as source cells, whose ligands bind to endothelial and AT2 epithelial cell receptors (**Figure 6B-right**). Monocytes predominantly interacted with other immune cells as both source or receiver cells, and the interaction numbers among immune cells were more sporadic compared to those between immune and non-immune cells (**Figure S5A**). These results suggest that monocytes may play a central role in orchestrating the immune microenvironment.

**Figure 6.**
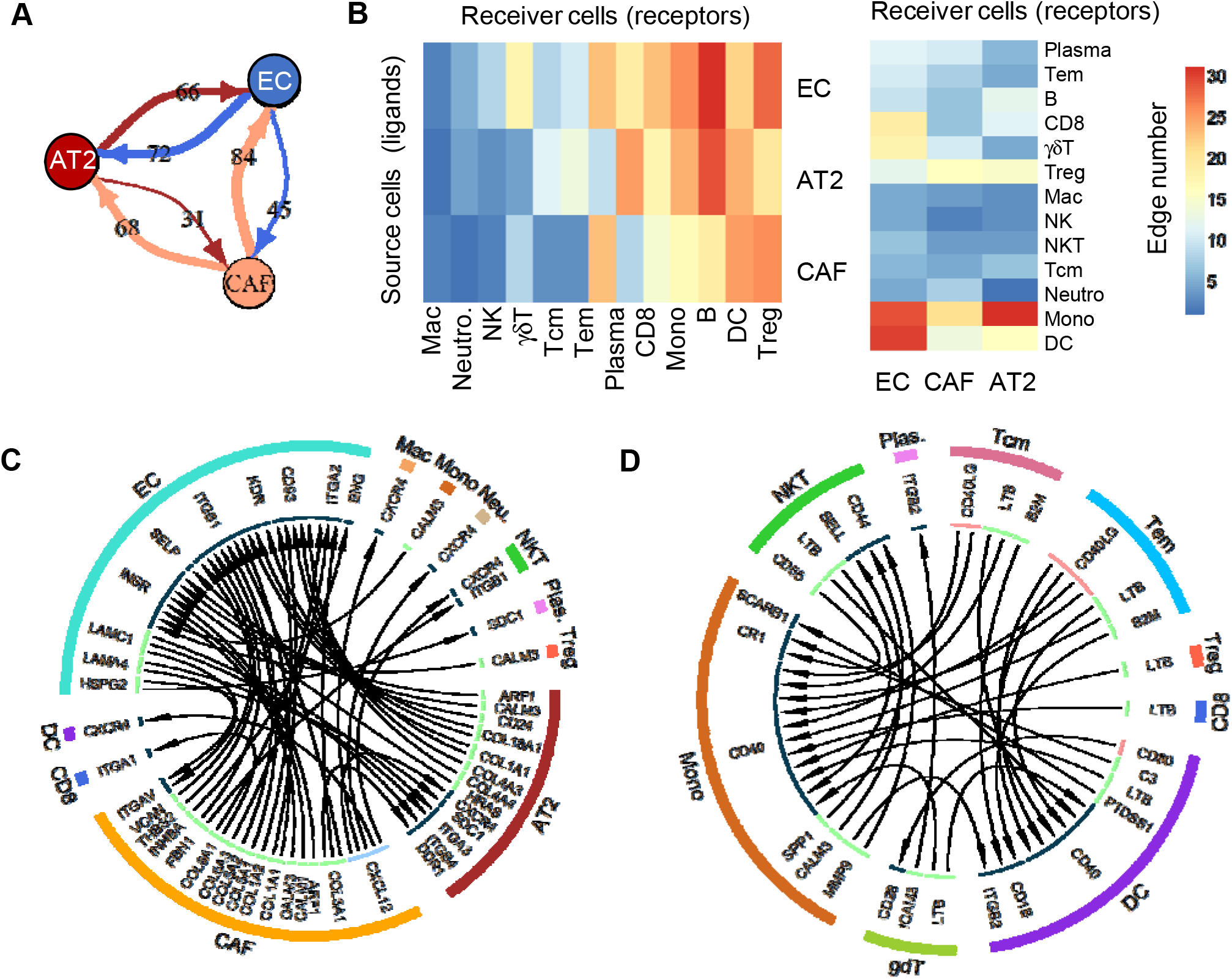
Interaction between cell types through ligand-receptor (LR) analyses. (A) Interaction among non-immune cells. Values on lines indicate the number of activated LR interactions in subsolid/solid lesions by comparing their scores to that of normal lung (B) Heatmap illustrating the number of unidirectional interactions from non-immune to immune (left) and vice versa (right) activated in subsolid/solid lesions compared to normal tissue. Rows indicate the source cells expressing ligand while columns represent receiver cells expressing receptor genes (C) Circos plot illustrating the unidirectional LR interactions between non-immune and immune cells activated in subsolid/solid lesions compared to normal lung due to the differentially expressed ligands and receptors in the indicated cell types. (D) Circos plot illustrating the significant LR interactions among immune cells due to the differentially expressed ligands and receptors in indicated cell types.

Two critical networks were identified, illustrating the interactions either between immune and non-immune cells or between immune cells, by assessing cancer-associated interactions of ligand and receptors identified as DEGs in related cell types. CAFs were identified as communicating with immune cells, including NKT, DC, macrophages, and neutrophils, by interaction with CXCR4 and the highly expressed CAF ligand *CXCL12* (**Figure 6C**). Potentially important communications between immune cells included an increase in interaction between myeloid cell receptor *CD40* (DC and monocytes) and its ligands (*CD40LG* and *LTB*) on T cells (NKT, Tcm, and Treg) in early ADC compared to the normal lung tissue (**Figure 6D**). These results suggest that CAF-induced alterations in the microenvironment may impact lesion immune infiltration. To understand the ligand-receptor interactions potentially contributing to B cell infiltration of subsolid lesions, potential B cell interactions were selected for further evaluation if their ligands were significantly up-regulated in malignant compared to normal tissues (FDR<0.01). Integrins (*ITGB1* and *ITGB4*) in B cells were found to interact with tip-like ECs having high expression of laminins (*LAMC1, LAMB1*, and *LAMA4*) as well as CAF collagens (*COL3A1, COL5A1*, and *COL6A3*) (**Figure S5B**). This suggests that the laminin- and collagen-enriched microenvironment created by cancer-associated endothelial cells and fibroblasts may attract B cells to lesion sites.

## Discussion

Recent studies of pulmonary premalignancy enhance the understanding of molecular and cellular characteristics in early spectrum lung cancer (Beane et al., 2019; Kadara et al., 2017; Teixeira et al., 2019). However, detailed single-cell level assessments of both the immune and non-immune contexture have not yet been applied to gain insights into the earliest events in the pathogenesis of lung ADC. In the present study, we provide an atlas of cellular components via single-cell transcriptome profiling of subsolid and solid lung ADC lesions, initially identified by clinical imaging. The key findings are: 1) a reduction in NK and NKT cells accompanied by an increase of CD4^+^ T, B cell, and DC populations in subsolid lesions as well as increased Treg cells. There were reduced effector T cells and cDC1 subtypes both of which are essential for anti-tumor immunity; 2) altered gene expression signatures in subsolid lesion-associated malignant alveolar type 2 pneumocytes, endothelial cells and fibroblasts promoting angiogenesis, inflammatory responses and extracellular matrix remodeling; and 3) modulation of immune cell infiltration by non-immune cells through ligand-receptor interactions, which ultimately may alter the microenvironment in subsolid lesions.

The numeric changes in immune cell populations observed in subsolid lesions were consistent with previous studies evaluating ADC solid lesions at a single-cell resolution (Lambrechts et al., 2018; Lavin et al., 2017), suggesting that the initial changes in the immune microenvironment may persist in advanced lung cancer. In the current study, correlates of immune cell functional activity were found at the transcriptome level. For example, tumor-infiltrating NKT cells exhibited decreased cytolytic capacity while expressing cytokines associated with iNKT2, which have the capacity to mitigate effective immune recognition. One of these cytokines, amphiregulin (AREG), engages the EGFR signaling pathway and can enhance immunosuppression by Treg cells, which in our study were found to be significantly elevated in subsolid lesions (Zaiss et al., 2013). AREG also has the capacity to directly stimulate KRAS mutant cancer cell proliferation and survival as well as circumventing tyrosine kinase inhibitor effectiveness in the context of *EGFR* mutant lung cancer therapy (Elangovan et al., 2018; Ishikawa et al., 2005; Taniguchi et al., 2017; Xu et al., 2019). The majority of our cohort’s malignant specimens harbored either *EGFR* or *KRAS* mutations (**Table S1**), thus suggesting that the upregulation of *AREG* by NKT cells could impact both disease progression and treatment outcomes.

A reduction in the correlates of infiltrating CD8^+^ T cell cytolytic function, indicated by the decreased expression of *IFNG* and *GZMB*, was observed in subsolid lesions. Although an increased percentage of CD4^+^ T cells was observed in subsolid lesions compared to normal tissue, we observed an increase in Treg cell percentage and a reduced Tem to Treg ratio. The two activated T cell clusters (clusters 14 and 21) predominantly originating from the specific subsolid lesions with both non-invasive and invasive histology, suggest that CD4^+^ T cells infiltrating early premalignant lesions may have distinct features from those in more advanced disease. Future studies will be required to define the characteristics of infiltrating lymphocytes associated with regressive or progressive premalignant lesions. The promising results from trials assessing neoadjuvant immunotherapy for NSCLC, together with the observed alterations in infiltrating lymphocyte functions in pulmonary premalignancy, suggest that immune-based therapies may be most effective in preventing tumor progression when administered at the earliest points of disease (Forde et al., 2018).

The diversity in DC subsets is evident in differences in developmental origin, phenotype, gene expression and function (Wculek et al., 2020). cDC1 and cDC2 subsets may play central roles in dictating the fate of the developing tumor. We observed increased infiltrating cDC2s in subsolid lesions, which correlated with increased CD4^+^ T and B cells. cDC2 have the capacity to interact with CD4^+^ T cells in the context of neoantigen presentation by MHC class II. Recent studies demonstrate that MHC II neoantigens and cDC2 are crucial for shaping an effective anti-tumor response (Alspach et al., 2019; Binnewies et al., 2019; Laoui et al., 2016). Among DC subsets, the cDC2 gene signature was the most strongly associated with a positive prognosis in lung ADC patients (Zilionis et al., 2019). We found an abundance of Treg cells in subsolid lesions, along with increased cDC2, emphasizing the importance of the documented capacity of Treg cells to impair the antitumor properties of cDC2 (Binnewies et al., 2019). The cDC1 subset possesses a heightened capacity to cross-present exogenous antigen and activate CD8^+^ T cells (Wculek et al., 2020). Recent studies highlight the role of NK cells in attracting cDC1 to the tumor site (Spranger et al., 2017). Via the production of *CXCL9* and *CXCL10*, intratumoral cDC1 are required for type 1 cytokine-secreting T cell effector trafficking to the tumor site. These *CXCR3* expressing T lymphocytes have the capacity for IFNγ production that enhances tumor MHC I expression as well as the continued heightened production of the IFNγ-inducible chemokines, *CXCL9* and *CXCL10*. Thus, the paucity of infiltrating NK cells may dictate the diminished cDC1 observed here in most subsolid lesions, potentially heightening the threat of disease progression. Studies suggest that the intratumoral injection of autologous gene-modified DC, termed *in situ* vaccination, may resolve this DC deficit (Yang et al., 2004). Early phase clinical trials of *in situ* vaccination in advanced NSCLC indicate induction of specific systemic antitumor immunity (Lee et al., 2017). Recent advances in the *in vitro* propagation of cDC1 and the *in situ* vaccination approach may make it feasible to evaluate these strategies for early interception of lung premalignancy (Lee et al., 2017; Perez and De Palma, 2019; Zhou et al., 2020).

Alterations in subsolid lesion non-immune cell profiles, including down-regulation of epithelial AT2 cell lineage markers, are consistent with the observations in organoid models in the context of *KRAS* activation (Dost et al., 2020). AT2 cells from subsolid lesions also exhibited deregulated gene expression and signaling pathways associated with tumor progression in the current study. Interrogation of subsolid ECs and CAFs revealed increased gene expression signatures consistent with the promotion of angiogenesis and extracellular matrix remodeling. A CAF subpopulation (cluster 9) was identified as exclusively present in malignant lesions. Cells in this subtype highly expressed the ligand *CXCL12*, whose receptor *CXCR4* was found to be upregulated both in AT2 and immune cells, including DCs, macrophages and NKT cells. Via CXCR4, CXCL12 stimulates increased tumor cell migration (Huang et al., 2007) as well as increased secretion of matrix metalloproteinases, mediating the degradation of the extracellular matrix (Ghosh et al., 2012). Targeting the CXCL12/CXCR4 axis was found to have synergistic effects with immune checkpoint blockade in murine models (Feig et al., 2013). Although it has been demonstrated that tumor intrinsic properties define distinct immune infiltration in the tumor microenvironment (Li et al., 2018; Skoulidis et al., 2015), our studies of the LR interactions between non-immune and immune cells reveal potential roles of tumor stroma elements, including CAFs and ECs, in shaping the infiltrating immune cell contexture. CAFs appear to be the predominant cell type modulating the interactions with other non-immune and immune cells in subsolid lesions. Therefore, interrogating CAFs may hold promise in identifying biomarkers as well as therapeutic targets for early detection and interception of lung cancer.

In summary, our results provide insights into alterations at the single-cell transcriptome level in both immune and non-immune cells in early spectrum lung ADC. Creating a premalignancy and early spectrum lung ADC atlas will enhance our understanding of disease pathogenesis and provide opportunities for early diagnosis and intervention.

## Acknowledgements

This work was supported by A Stand Up to Cancer-LUNGevity-American Lung Association Lung Cancer Interception Dream Team Translational Cancer Research Grant (Grant Number: SU2C-AACR-DT23-17 to S.M. Dubinett and A.E. Spira). Stand Up to Cancer is a division of the Entertainment Industry Foundation. Research grants were administered by the American Association for Cancer Research, the scientific partner of SU2C, NCI HTAN PCA 1U2CCA233238, NIH/NCI MCL 1U01CA196408, NIH/NCI EDRN 1U01CA214182, Merit Review and SDR Research funding from the Department of Veterans Affairs (S.M.D), TRDRP 27IR-0036 (K.K.), Thoracic Surgery Foundation research grant (J.Y.). This work also received support from Lung PCA Consortium and HTAN Consortium (https://humantumoratlas.org).

---

**Figure S1.**
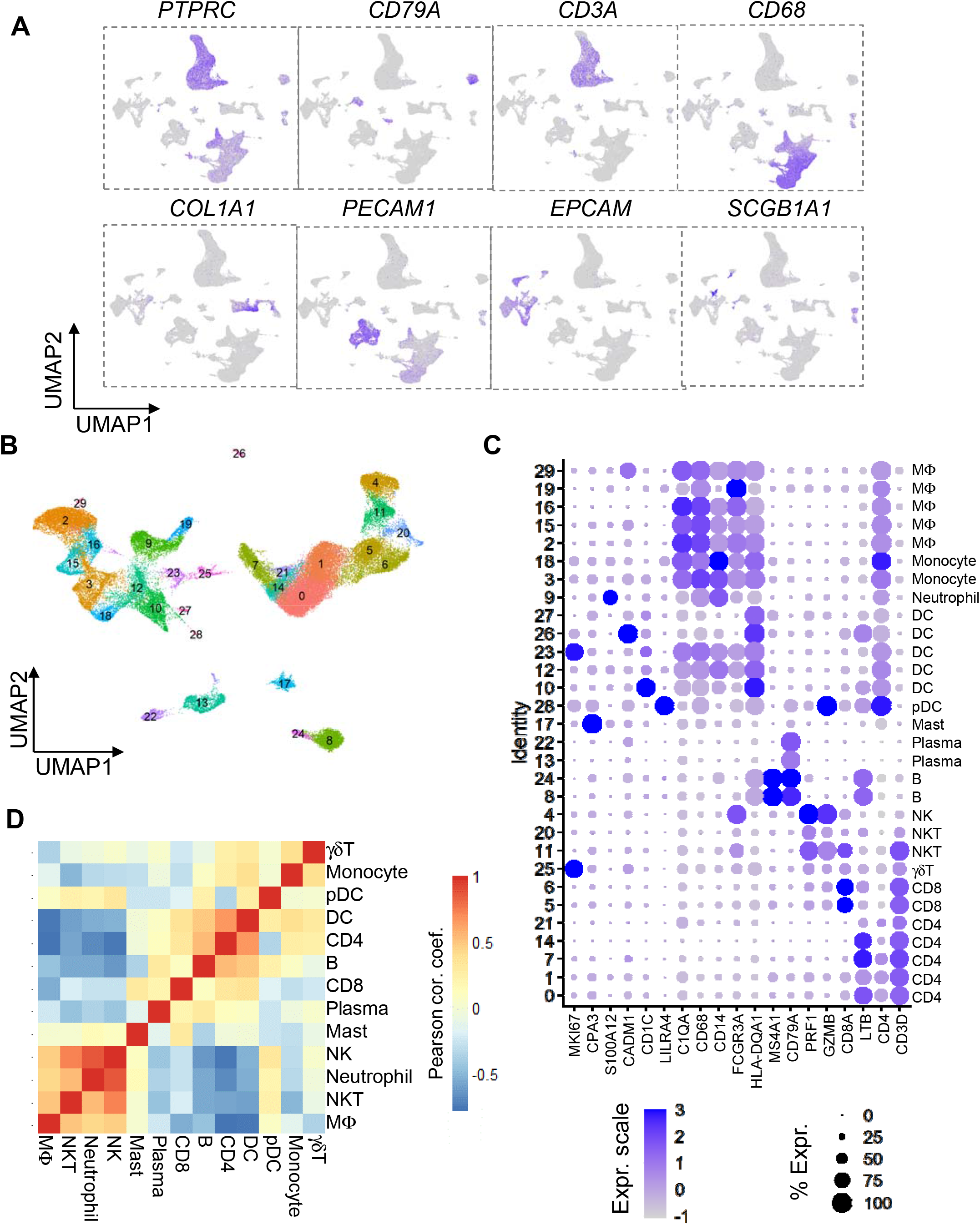
Characterization of immune cell types based on lineage markers. (A) Single-cell gene expression of representative markers for: immune cells (CD45, *PTPRC*) and their sub-lineage cells including B cells (*CD79A*), T cells (*CD3A*), myeloid (*CD68*), as well as fibroblast (*COL1A1*), endothelial (*PECAM1*), epithelial (*EPCAM*), and secretory cells (*SCGB1A1*) (B) UMAP plot visualizing 30 clusters formed by 60,509 immune cells (C) Bubble plots illustrating expression of lineage markers in individual clusters. The bubble size represents the percentage of cells in the cluster expressing (UMI > 0) the indicated gene, while the colors represent the average expression in the cluster. The major cell lineages are natural killer (NK), natural killer T (NKT), CD8^+^ T (CD8), CD4^+^ T (CD4), gamma delta T (γδT), B, plasma mast, dendritic cell (DC), plasmacytoid DC (pDC), macrophage (MΦ), monocyte (Mono) and neutrophil (Neutro) (D) Heatmap indicating the correlation between immune cells based on their relative abundance.

**Figure S2.**
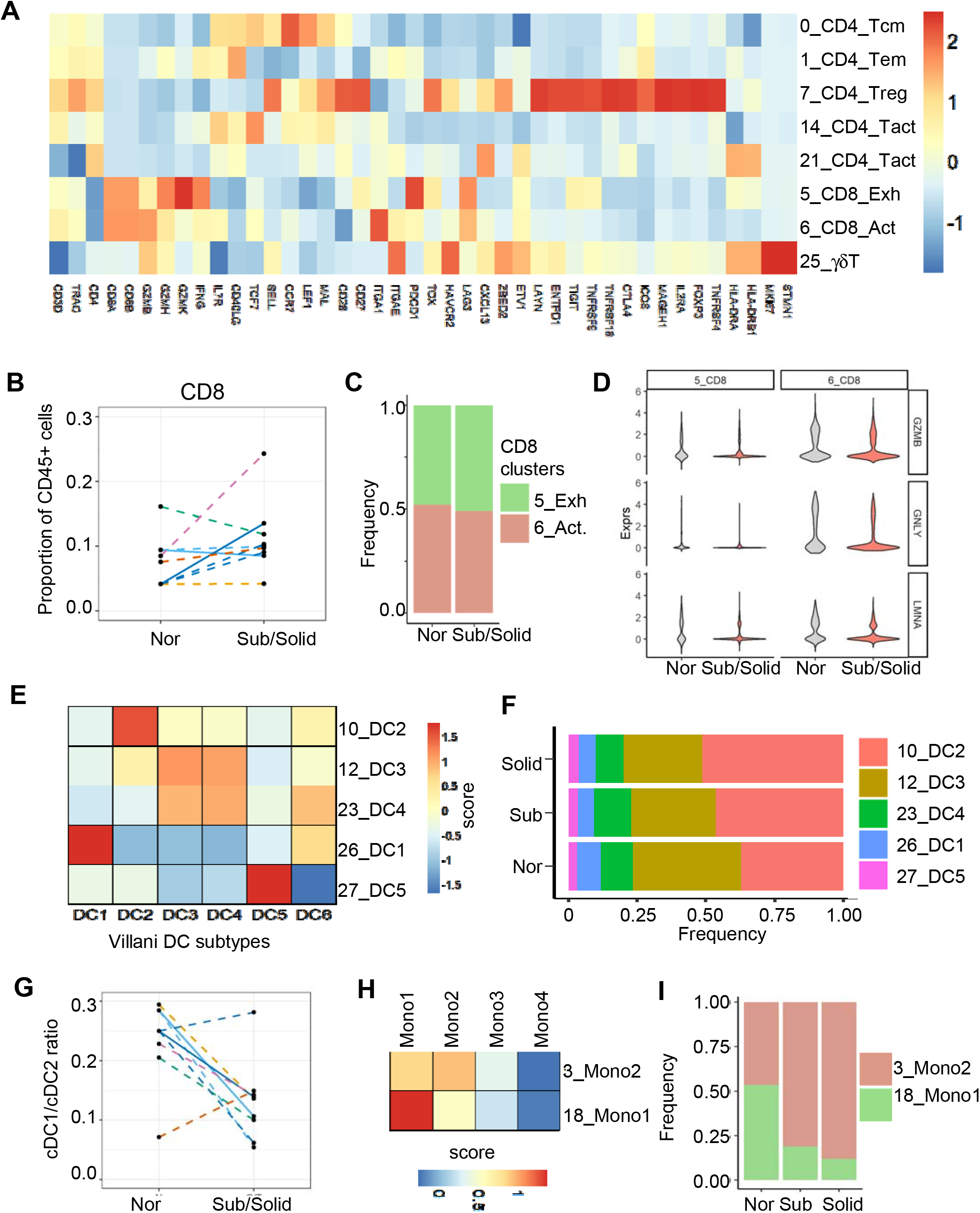
Functional annotation of immune cell sub-lineages. (A) Heatmap representing expression of known T cell markers in clusters annotated as T cells, including CD4^+^, CD8^+^, and gamma delta T cells (γδT) (B) Line plot illustrating CD8^+^ T cell frequency between normal and abnormal lesions. Line colors represent individual patients, while the line pattern indicates the normal-subsolid (dashed) and normal-tumor (solid) relationship (C) Distribution of CD8^+^ T cell subtypes in different lesion types. Colors indicate subtypes (D) Violin plots illustrating gene expression of DEGs between malignant- and normal-derived CD8^+^ T cells (E) Heatmap (right) representing scores of indicated DC gene modules (columns) in the related DC-associated clusters (rows). (F) Distribution of DC subtypes in different lesion types. Colors indicate subtypes (G) Reduction of cDC1:cDC2 ratio from normal to associated subsolid and solid lesions. Line colors represent individual patients, while the line pattern indicates the normal-subsolid (dashed) and normal-tumor (solid) relationship (H) Heatmap (right) representing scores of indicated monocyte gene modules (columns) in the related monocyte-associated clusters (rows) (I) Distribution of DC subtypes in different lesion types. Colors indicate subtypes

**Figure S3.**
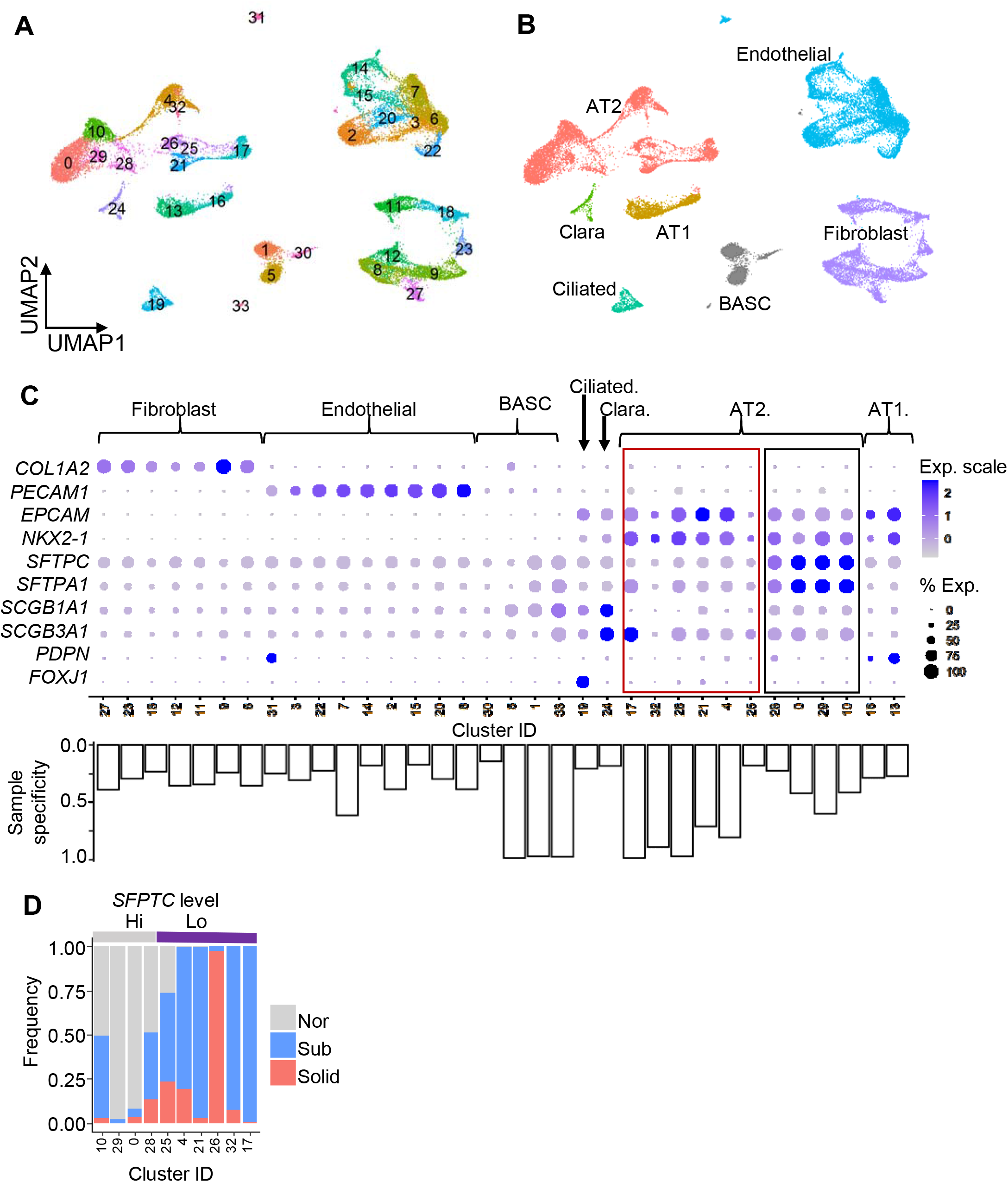
Characterization of non-immune cells. (A) UMAP plot visualizing 34 clusters formed by 28,129 non-immune cells. Colors represent clusters (B) Annotated non-immune clusters. Colors represent cell types (C) Bubble plots (top) illustrating canonical gene expression. Seven cell types, including bronchoalveolar stem cells (BASCs), were captured. Boxes indicate two AT2 groups having low (red) and high (black) *SFTPC* expression levels. The bar plots (bottom) represent sample specificity defined as the maximal percentage of a sample’s contribution to the indicated clusters. (D) Lesion-based contribution to the clusters classified according to SFTPC expression levels Colors indicate lesion types

**Figure S4.**
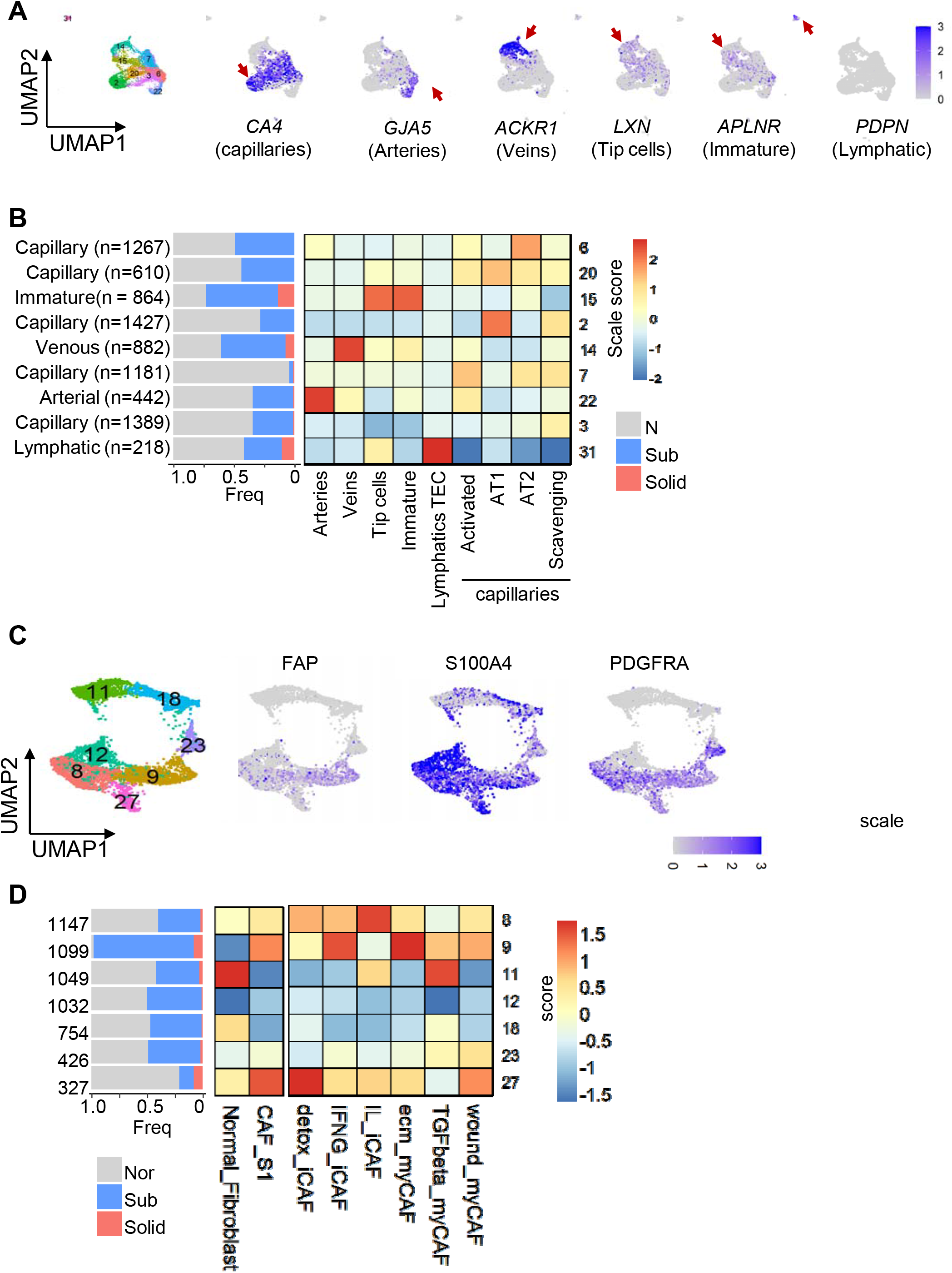
Characterization of ECs and fibroblasts. (A) Expression of canonical markers associated with endothelial sub-lineages (indicated in parentheses) (B) Lesion-based contributions (left) to clusters annotated as ECs. Heatmap (right) representing scores of indicated gene modules (columns) in clusters (rows). (C) Expression of known CAF markers in clusters annotated as fibroblasts (D) Lesion-based contributions (left) to clusters annotated as fibroblasts. Heatmap (right) represents scores of indicated gene modules (columns) in clusters (rows).

**Figure S5.**
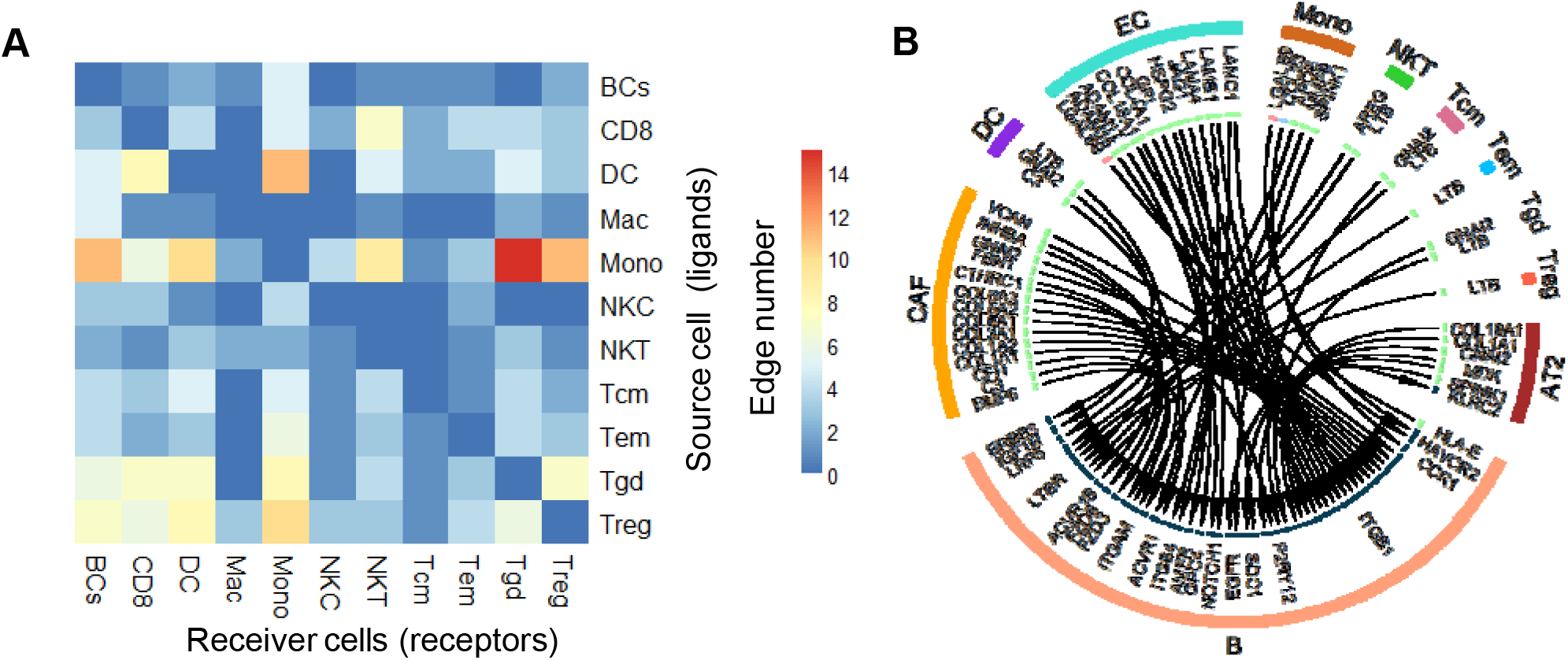
LR interactions among immune cells. (A) Heatmap illustrating the number of significant LR interactions among immune cells. Rows indicate the source cells expressing ligand, while columns represent receiver cells expressing receptor genes (B) Circos plot illustrating the interactions from other cell types to B cells

## Supplemental Tables

**Table S1. Clinical and radiological characteristics of subsolid and solid lesions**

**Table S2. DEGs in Cluster 20 NKT cells between subsolid/solid lesions and non-involved tissue**

**Table S3. DEGs in AT2 cells between subsolid/solid lesions and non-involved**

## STAR Methods

### RESOURCE AVAILABILITY

#### Lead Contact

Further information and requests for resources and reagents should be directed to the Lead Contact, Steven M. Dubinett

#### Data and Code Availability

The dataset generated in this study is available in Human Tumor Atlas Network – Data Coordinating Center (https://humantumoratlas.org/htan-dcc/). The raw RNA sequence is under restricted access due to patient privacy concerns.

### EXPERIMENTAL MODEL AND SUBJECT DETAILS

#### Human Specimens

Fresh non-involved normal lung and disease tissues were obtained from patients undergoing lung cancer surgery at UCLA Medical Centers (Santa Monica and Los Angeles, CA) after obtaining written informed consent. The study and its protocols were reviewed and conducted with the approval UCLA IRB (IRB # 10-001096). Once the lung resection was performed, the tissue was sent immediately and fresh to pathology. The solid and subsolid lesions were identified by the operating thoracic surgeon, based on careful palpation and correlation with the imaging. In cases where the diagnosis had not already been verified by needle biopsy, frozen section analysis with review by a dedicated thoracic pathologist confirmed early spectrum lung adenocarcinoma. A portion of the primary subsolid lesion, any associated solid tumor, as well as a sample of normal lung tissue from a grossly normal site of the lobectomy specimen, distant (> 2cm) from the tumor site(s), was then harvested. Mutation status was assessed by the targeted DNA sequencing and FISH per clinical protocol. PD-L1 was assessed by immunohistochemistry.

#### Patient Clinical Characteristics

Samples were collected from 6 patients whose clinical features were described in Table S1.

### METHOD DETAILS

#### Sample dissociation for scRNA-Seq

Resected tissues were placed on ice in RPMI medium immediately after resection and delivered to the lab for tissue dissociation. The single cell dissociation protocol was adapted from Leelatian et al. (Leelatian et al., 2017). In brief, dissociation was performed in RPMI medium supplemented with 10% FBS. Tissues were sliced to approximately 1mm^3^ pieces and dissociated in 200 Units/ml collagenase type II (Sigma Aldrich, #C6885) and 100 Kunitz Units/ml DNAse I (Sigma Aldrich, #DN25) at 37°C for approximately 1 hour until homogeneity followed by passing through a 40 μm strainer to remove cell aggregates and red blood cell lysis with 1 ml of ACK buffer (Sigma Aldrich, #11814389001). Cells were resuspended in 5 ml DPBS + 0.04% BSA, counted, and immediately used to prepare the sequencing libraries.

#### Single-cell RNA sequencing (scRNA-Seq) and read alignment

The 10X Genomics platform (10X Genomics, Pleasanton, CA) was utilized for assessing human single-cell transcriptome. Single-cell encapsulation, library construction, and sequencing were performed at Technology Center for Genomics and Bioinformatics at UCLA according to the manufacturer’s protocols. The Chromium™ Single Cell 3’ Library & Gel Bead Kit v2 and v3 were used for library preparation. Libraries were sequenced utilizing Illumina NovaSeq 6000 instrument. CellRanger 3.1.0 software (10X Genomics) was utilized to align and annotate reads based on human genome assembly GRCh38p13 and gene annotation GENCODE34 and then generate count matrices.

#### Single-cell data filtering, normalization, and batch adjustment

Count matrices generated by CellRanger 3.1.0 were processed by following Seurat pipeline (Stuart et al., 2019). In brief, count matrices of individual samples were first combined and filtered out low-quality cells, which had > 17% mitochondrial content and < 475 detected genes. A total of 88,638 cells were retained for further analysis. The data was normalized and batch-adjusted following the Seurat Standard Workflow pipeline in which the top 2000 highest variance genes were utilized to find anchors and integrate data from different batches.

#### Single-cell cluster analysis and annotation

We performed cell clustering on the batch-adjusted data to separate immune cells from non-immune cells *in-silico* by utilizing the top 50 principal components to determine the k-nearest neighbors of each cell and visualize cells by UMAP. The Louvain algorithm-based cluster identification was tested at a range of resolution from 0.2 to 1.5 to determine if the increase of resolution value produced new clusters associated with a specific sample. We decided to utilize the optimized resolution = 1.0, which was high enough to obtain clusters associated with cell lineage identity and still minimize the number of clusters associated with a specific sample. The separated immune and non-immune cells were then reanalyzed independently to determine the highest variance genes, re-calculate principal components, and cluster and visualize cells by utilizing the top 30 new calculated principal components.

We annotated obtained cell clusters by a 2-tier pipeline. We first curated cell lineage markers from multiple databases, including PanglaoDB (Franzen et al., 2019) and CIBERSORT (Newman et al., 2015), and previously published articles, in which single-cell transcriptomes were available for the particular immune lineages, such as Villani et al. (Villani et al., 2017) for dendritic cells and monocytes, Zillionis et al. (Zilionis et al., 2019) for macrophages and neutrophiles, Leader et al. (Leader et al., 2020) for T cells. We then perform the following pipeline:

1. In tier 1 of the pipeline, we utilized the systematic approach to determine if the lineage markers of a cell type enriched as the positive marker genes in a cluster. The enrichment score of cluster *C* as cell type *T, R_C,T_*, was calculated as:

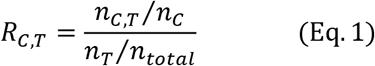

where *n_C,T_* is the number of genes that identified as positively expressed markers in both the cluster *C* and cell type *T*, while *n_c_, n_T_* and *n_total_* are the total numbers of positively expressed markers in cluster *C*, cell type *T*, and the total number of genes in detected by scRNA-seq, respectively. The positively expressed markers in each cluster were identified by *FindAllMarkes* utilizing the MAST approach.
2. For each cluster, we ranked its enrichment scores associated with different cell types from high to low. If the deviation between 1^st^ and 2^nd^ highest scores was > 1, the cell type with the highest score was assigned to the cluster. If not, we performed the next analysis step based on the expression of canonical markers
3. For each cluster, we compared canonical expressions of candidate cell types predicted from the previous step. The cluster was assigned to the cell lineage with the higher average expression of canonical markers.

#### Annotating functional subtypes by known gene modules

We observed some cell lineages composed of multiple clusters were associated with their subtypes or biological stages. We determined their functional subtypes based on their literature signature gene modules. In brief, we first modified the existing gene modules to maintain only the mutually exclusive genes among them. Each signature gene set has its module score determined as the z-score average across all module genes for each cell. The module score at the cluster level was the average across all cells in the cluster.

#### Analysis of differentially expressed genes

Two different approaches, pseudo-bulk edgeR (Robinson et al., 2010) and single-cell model MAST (Finak et al., 2015), were utilized to determine differentially expressed genes (DEGs) between subsolid/solid and normal AT2 cells. The final DEGs were the intersection of both approaches. In brief, we utilized raw read counts of cells identified as AT2 cells (regardless of their *SFTPC* expression level) in each approach, so that we added a variable incorporating patient identity besides tissue histology when modeling expression changes. A DEG was identified if its fold-change > 2 and FDR < 0.1 in pseudo-bulk edgeR approach, while it required the coefficient > 0.5 and FDR < 1e-50 in MAST approach.

The DEGs between subsolid/solid lesions and normal tissue in cell types other than AT2 were assessed by utilizing the MAST approach since their transcriptome profiles were usually more homogeneous among samples than the AT2 population. Furthermore, we excluded cells in clusters associated with a specific sample (>70% cells in the cluster were from a sample) in analyzing the DEGs in these cell types.

#### Gene set enrichment analysis

We used Fisher’s exact test to determine the pathways or biological processes enriched by DEGs, and adjusted p-values were with the Benjamini-Hochberg method for multiple hypothesis testing. The statistically significant altered pathways (FDR <0.05) were then re-analyzed by the ranked-based GSEA approach (Subramanian et al., 2005) utilizing the Bioconductor *fgsea* package (Sergushichev, 2016) to assess the direction of deregulation in the subsolid/solid lesion samples. We utilize gene sets obtained from the Molecular Signature Database (MsigDB) version 7.1.

#### Ligand-receptor interaction and cell-to-cell afinity among cell types

To evaluate the communication between cell types through their ligand-receptor interactions, we defined the strength of each interaction of given ligand *l_i_* in cell type A to its related receptor *r_i_* in cell type B as the product of average ligand expression across all cells of cell type A and average ligand expression across all cells of cell type A as suggested by Kumar et al (Kumar et al., 2018):

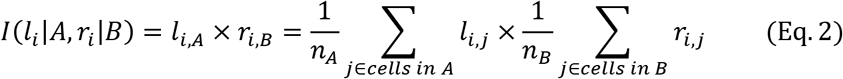

where *n_A_* and *n_B_* are the total number of cells in cell type A and B, respectively, and *l_i,j_*(*r_ij_*) is the expression (in log scale) of gene *l_i_* (*r_i_*) in cell *j*. For a given cell type A and B, an LR pair would have two different represented scores calculated by assuming the ligand in A interacting with the receptor in B, and vice versa. Furthermore, interaction scores were also computed for each lesion since lesions might have different microenvironments even though they are from the same subject. We utilized the LR database in the R-based *iTALK* package (Wang et al., 2019), which categorized literature LR pairs into four categories: immune checkpoint, cytokine/chemokine, growth factors, and others, based on ligand functions. To calculate average gene expression of each cell type, we selected cells in clusters that were (1) composed of cells from multiple samples, and (2) associated with disease status. For instance, we selected endothelial cells in cluster 2 (normal), 14 and 15 (malignancy), and fibroblast cells in cluster 11 and 18 (normal), and 8 and 9 (malignancy).

We performed two-sided Wilcoxon rank-sum tests comparing scores between malignant and normal samples for each specific LR interaction. A specific LR interaction was identified as significance in subsolid/solid lesion compared to normal if (1) its Wilcoxon rank-sum p-value < 0.05, (2) fold change of their scores between malignant and normal samples > 2 and (3) either ligand (*l_i,A_*) or receptor (*r_i,B_*) expression was differentially expressed (FDR<0.1) in malignant lesion compared to normal lung tissue.

### QUANTIFICATION AND STATISTICAL ANALYSIS

Statistical testing was performed utilizing R 3.6. We utilized the linear mixed-effects model (R *lmerTest* package) to incorporate individual patient variation in evaluating changes between groups. To eliminate the bias due to small sample size, data from solid lesions were not included in the statistical tests at sample level, but at cell level analysis. Appropriate rank-based statistical tests were applied according to the nature of variables. The tests used to determine statistical significance were quoted next to the p-values and in the appropriate figure legends.

